# Enhanced neural encoding of individual exemplars in the Super-Recogniser brain is specific to faces

**DOI:** 10.64898/2026.07.20.739570

**Authors:** Martina Ventura, Tijl Grootswagers, Manuel Varlet, James Dunn, David White, Genevieve L. Quek

## Abstract

Super-Recognisers are known for their exceptional ability to encode and remember faces. Recent work has found robust neural codes for face identity in brain activity. Here we test whether the enhanced individuation of face identity in super-recogniser neural codes extends to exemplars from non-face categories. We recorded 64-channel electroencephalography (EEG) as 22 Super-Recognisers and 20 typical recognisers viewed rapid, randomised sequences of faces, dogs, houses, and cars (10 different images of 10 different exemplars per category). Using a neural decoding approach, we found that Super-Recognisers’ neural responses contained stronger and more temporally sustained information discriminating individual face identities. Moreover, Super-Recognisers displayed increased inter-individual consistency in the organisation of their face-identity representations, suggesting a more stable and/or canonical representational structure in individuals with superior face recognition abilities. Crucially, there was no evidence to suggest that these encoding advantages generalised to non-face categories: neural discrimination of individual dogs, cars, and houses was comparable across groups, and group-differences in inter-subject consistency were evident only for faces. Together, these findings suggest that enhanced perceptual encoding underlying exceptional face recognition is highly category-specific, supporting the view that expertise in this group reflects specialised mechanisms for processing facial identity rather than broadly enhanced visual discrimination capacities.

## 1. Introduction

Perceptual and cognitive phenomena can vary widely across individuals, reflecting meaningful differences in how sensory information is processed, encoded, and remembered. Inter-individual variability is particularly prominent in the everyday phenomenon of human face recognition: Although most of us successfully recognise faces of people we know without issue, a subset of individuals encode and remember face identities from relatively little input, far exceeding the capabilities of the average person (Russell et al., 2009).

So-called ‘Super-Recognisers’ have been a key focus of research aimed at understanding the mechanisms underlying individual differences in face recognition and perceptual expertise more broadly. To date, their abilities have largely been characterised via tasks involving face-identity, with some less pronounced advantages in non-face (object) recognition also reported (Bobak et al., 2016; Dunn et al., 2023). This has led to the assumption that Super-Recognisers’ abilities reflect perceptual and cognitive mechanisms that are at least partly face-specific (Davis et al., 2016; Bennet & Bate., 2017), reflecting a wealth of neuropsychological evidence showing dissociations between neural systems for face versus generic object recognition (Gerlach & Starrfelt, 2024; Duchaine & Nakayama, 2005; McMullen et al., 2000). Studies of individual differences in the broader population also point to these abilities being at least partly dissociable (White & Burton, 2022), possibly underpinned by independent genetic contributions (Shakeshaft & Plomin, 2015).

Recent findings have challenged this position, raising the possibility that Super-Recognisers’ abilities may extend beyond strictly face-selective mechanisms (Hendel et al., 2019; Nador et al., 2025; Dunn et al., 2023; Towler et al; 2023). In particular, a recent comprehensive within-subject assessment of super-recognisers’ cognitive capabilities by Stehr et al. (2026) suggests they enjoy robust advantages over typical observers not only on classic face memory tasks (e.g., CFMT+; Duchaine & Nakayama, 2006), but also on non-memory-related face processing tasks, and non face-related longer term memory tasks involving visual and auditory stimuli. Crucially, these advantages did not extend uniformly across *all* cognitive domains, with more modest and inconsistent effects observed for tasks involving neither faces nor longer-term memory demands (Stehr et al., 2026).

Characterising the mechanisms that give rise to this pattern of performance is a clear next step in understanding individual differences in visual expertise. Yet behavioural performance measures alone have limited potential to reveal how domain-general and face-specific mechanisms intersect, as standard tasks necessarily collapse multiple cognitive operations and stages of processing into a single outcome: An individual’s recognition performance can reflect not only consolidation, retrieval, and decision-related processes, but also differences in the initial perceptual encoding of visual information (Karuza et al., 2014; Ortu & Vaidya, 2017).

In contrast, time-resolved neuroimaging (e.g., electroencephalography - EEG) is well-positioned to tease apart effects arising during perceptual encoding and later retrieval/decision related processing, by examining the structure and temporal dynamics of neural responses elicited during passive stimulus viewing. To date, however, such investigations with Super-Recognisers have been extremely rare (Faghel-Soubeyrand et al., 2024; Nador et al., 2025). Interestingly, these studies have pointed to neural distinctions between Super-Recognisers and typical viewers that extend beyond strictly face-selective processes. Using EEG decoding and representational modelling, Faghel-Soubeyrand et al. (2024) showed that Super-Recogniser group membership could be predicted from neural responses to both face and non-face stimuli, with neural responses in this group displaying stronger links to both mid-level visual representations and later semantic representations. Similarly, Nador et al. (2025) have reported that where selective neural responses for face-identity and face-category are largely comparable in Super-Recognisers and typical viewers, general visual processing responses appear to be stronger in the former group.

Where these studies seem to allude to a link between superior face recognition ability and enhanced *domain-general* processing across multiple stages of visual and semantic representation, we recently provided compelling evidence that neural encoding of individual face identities is enhanced in Super-Recognisers relative to typical viewers (Ventura et al., 2026). Super-Recognisers brain activity implied a representational geometry for facial identity than better discriminated between individual faces compared to control participants, even in the absence of any explicit identity-related task. Moreover, the geometry of ‘facespace’ representations (Valentine, 1991), were more consistent across individual super-recognisers that for the comparison group, pointing to perceptual processing systems tuned to face identity. Although the timcourse of this difference pointed to higher-level object processing rather than low level processing differences, we were unable to confirm that these representational differences were face-specific.

Whether Super-Recognisers’ observed enhanced neural encoding of facial identity is underpinned by a general mechanism that enables improved discrimination of entities within any visual category is unresolved. Given that evidence suggests that highly discriminable perceptual representations during early visual processing can increase the stability and distinctiveness of information entering downstream mnemonic systems (Ye et al., 2024), if Super-Recognisers’ show stronger neural exemplar differentiation for non-face categories as well as faces, this would constitute a generally enhanced perceptual encoding mechanism in this group (Nador et al., 2021) that could account for their demonstrated advantage on memory tasks involving both face and non-face stimuli (Stehr et al., 2026).

Here we aimed to resolve this question using neural decoding of EEG data (Grootswagers et al., 2017) to characterize the perceptual encoding of individual faces, cars, houses, and dogs. Super-Recognisers (n=22) and typical recognisers (N=20) saw rapid randomised sequences containing 10 different images of 10 different exemplars from the four visual categories. The fast presentation rate ensured that participants could not visually explore the images, enabling us to capture neural representations of individual entities within each category in the absence of strategic memorisation or visual exploration. We found that neural representations of individual face identities were stronger and more temporally sustained in Super-Recognisers than in typical recognisers. Moreover, superior face recognition ability was associated with greater consistency in the organisation of face-identity representations across individual brains, suggesting a more stable and/or canonical representational structure in Super-Recognisers’ face encoding. Importantly, this advantage did not generalise to non-face categories: neural discrimination of individual dogs, cars, and houses was comparable across groups, providing no evidence for a domain-general enhancement of exemplar-level visual encoding. Together, these findings suggest that the enhanced perceptual encoding underlying exceptional face recognition is category-specific, supporting the view that expertise in this group reflects specialised mechanisms for processing facial identity rather than broadly enhanced visual discrimination capacities.

## 2. Methods

### 2.1 Participants

We recruited 42 participants consisting of 22 Super-Recognisers (15 F, *M*age = 31.44 years, *SD* = 5.16) and 20 typical recognisers (15 F, *M*age = 40.19 years, *SD* = 9.18 years). Participants were compensated monetarily for their participation. Super-Recognisers were sourced from an existing database of individuals who have previously completed a set of standardized face recognition tests. Here, we defined Super-Recognisers as individuals whose composite z score exceeded 1.7 standard deviations above the standardized mean on The Cambridge Face Memory Test–Long Form (CFMT+; Duchaine & Nakayama, 2006), the Glasgow Face Matching Test–Short Form (GFMT; Burton et al., 2010), and the UNSW Face Test (Dunn et al., 2020). typical recognisers were recruited from the general population and were screened in advance of in-person testing using the Prosopagnosia-Index 20 (Shah et al., 2015), to exclude participants with self-reported face recognition difficulties. After, to ensure comparability with the Super-recognisers sample, and to ensure they did not meet the Super-recogniser criteria, typical recognisers underwent the same battery of behavioural face recognition assessments as the Super-recognisers (i.e., CFMT+, GFMT, UNSW Face Test). This study was approved by the Human Research Ethics Committee at Western Sydney University (H15873) and conducted in accordance with the National Statement on Ethical Conduct in Human Research.

### 2.2 Procedure & Stimuli

The main experiment combined 64-channel EEG recording (BioSemi) with a rapid serial visual presentation (RSVP) paradigm. This approach leverages the capacity of the visual system to sustain multiple object representations in rapid succession and has been repeatedly shown to support reliable time-resolved multivariate decoding of visual representations, even when neural responses to successive stimuli partially overlap (Grootswagers et al., 2019; Robinson et al., 2019). Participants saw a 2.5 Hz image sequence containing images from four distinct categories: cars, dogs, houses, and faces. Each category contained 10 unique exemplars (e.g., *dog1, dog2, dog3*…), each of which was represented by 10 unique images (e.g., *dog1_img1, dog1_img2, dog1_img3*…), for a total of 40 images per category, and 400 images in the full stimulus set.

Stimuli were images sourced from the internet, selected to contain natural variability in facial appearance that define the computational challenge of face recognition in daily life. The images were presented at the centre of the screen on a 24-inch ViewPixx monitor. The presentation of stimuli and task control was managed using Python and PsychoPy (Peirce et al., 2019). The experiment consisted of 16 blocks, with each block containing all the 400 images presented in a randomized order. Images appeared at a rate of exactly 2.5 Hz, with each image presented for 250ms and followed by a blank inter-stimulus interval of 100ms (see Figure 1a). While this rapid presentation rate means that neural responses to successive stimuli inevitably overlap in time, randomised presentation order within each block ensures that responses to subsequent stimuli (N+1, N+2) cannot be systematically related to the target stimulus (N). Consequently, any condition-specific information decodable can be attributed to the target stimulus rather than to overlapping activity from subsequent images. This logic is well established in RSVP paradigms demonstrating reliable time-resolved decoding under conditions of temporal overlap (e.g., Grootswagers et al., 2019, 2024; Robinson et al., 2019).

**Figure 1.**
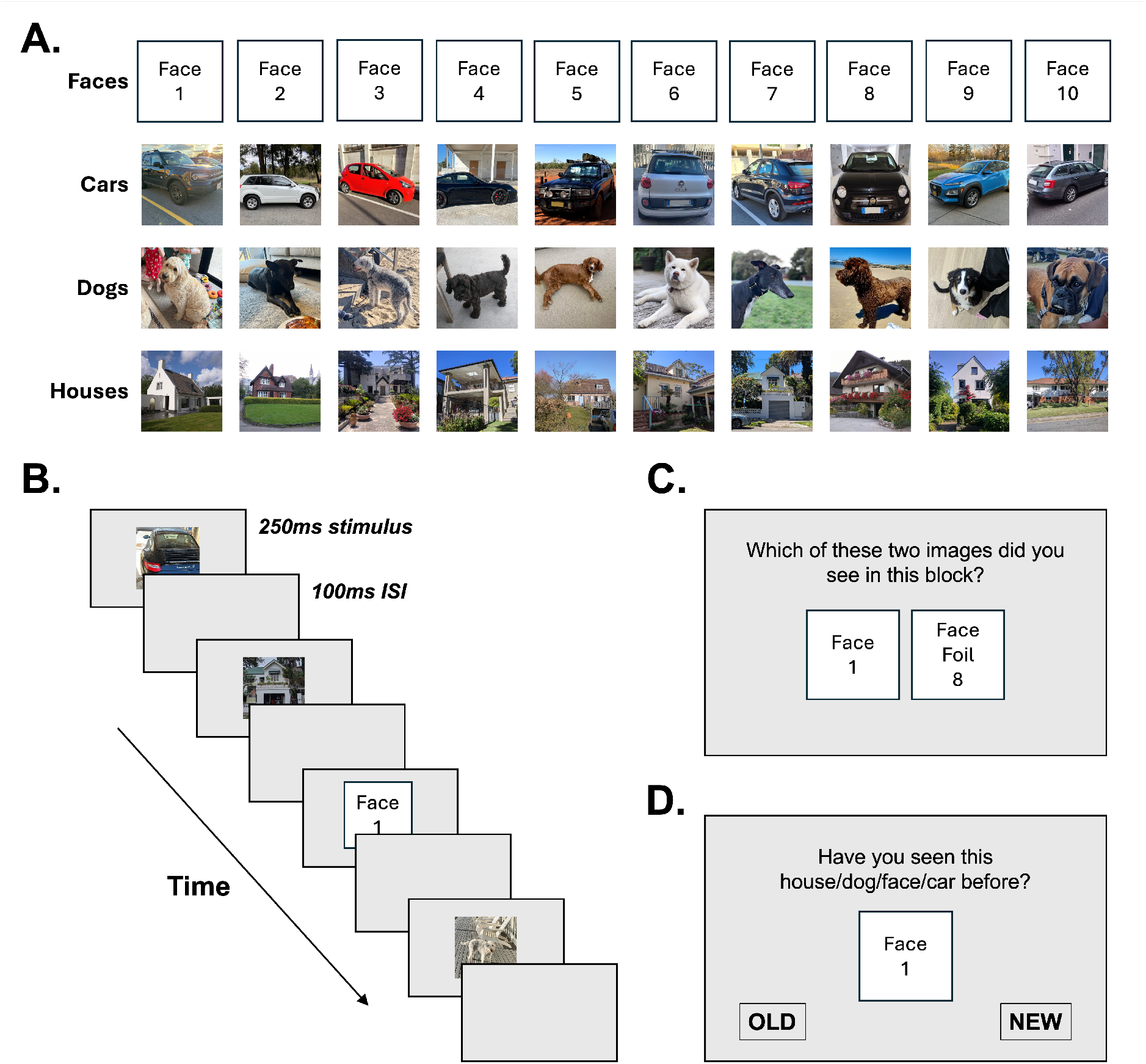
Experimental design. (A) We used 10 exemplars for each of the four visual categories. Each exemplar was represented by a set of 10 highly variable images, yielding 400 images in total. (B) In the main experiment, we recorded EEG as participants viewed the images in a 2.5 Hz RSVP-style stream (250ms stimulus duration + 100ms inter-stimulus interval, ISI). (C) After each sequence, participants completed a single attention check trial, selected which of two images had appeared in the preceding sequence. The target and foil were always drawn from the same category. (D) An example trial from the surprise memory task we included after the main experiment. Participants had to indicate if a depicted exemplar was previously encountered during the main experiment and were instructed to consider their familiarity with the *exemplar* rather than the exact image.

To encourage participants to maintain attention to the image sequence, we included a single memory trial following each sequence in which participants saw one randomly selected image from the block they had just completed and one foil that matched the category of the target (Figure 1C). Participants used the left and right arrow keys to indicate which of the two images had been in the just-seen sequence. The categories (i.e., cars, dogs, houses and faces) in the memory trials were randomized and counterbalanced.

Immediately following the main experiment, participants completed a short surprise memory test that was designed to verify that they had paid attention to the images presented during the main task. On each trial, participants saw a single image for 250 ms and then pressed the left or right arrow to indicate whether that exemplar was “New” or “Old” (Figure 1D). There were 160 trials: 40 images that had been presented during the main experiment (i.e., ‘seen-old’ - 10 per category), 40 new images of specific exemplars that had appeared in the main experiment (i.e., ‘novel-old’, 10 per category), and 80 foil images of never-before-encountered exemplars who had not appeared in the main experiment (i.e., 20 per category). There was no time constraint for the response; we emphasized to participants that they should consider whether the *exemplar* in the image was one they had encountered before, rather than the exact image. Based on prior literature (e.g., Stehr et al., 2026) we expected Super-Recognisers to outperform controls on this recognition task.

To ensure that the surprise memory test used previously-seen images that were most representative of their respective exemplars, we identified which of the 10 images for each exemplar should serve as the recognition target using visual similarity scores derived from DreamSim (Fu et al., 2023). DreamSim is a Deep Neural Network (DNN) adept at capturing hierarchical features in images, making it suitable for tasks requiring nuanced visual discrimination. The model computes a similarity score between pairs of images based on their feature representations, generating a distance metric that quantifies overall visual similarity between paired images. For each of the 40 exemplars used in the encoding phase, we calculated a single within-exemplar similarity score for each image by averaging across its similarity scores with the other nine images of that exemplar. This result reflects how similar that image was, on average, to the other images of the same exemplar. For each exemplar, the image whose within-exemplar similarity score was closest to the overall mean similarity of that exemplar was selected for inclusion as a previously-seen target image. This procedure ensured that the selected image was neither unusually similar nor dissimilar to the other images of that exemplar, but rather best represented the central visual characteristics of that exemplar.

### 2.3 EEG recording and pre-processing

EEG data were recorded using a 64-channel BioSemi Active-Two electrode system with a sampling rate of 2048 Hz (BioSemi, Amsterdam, The Netherlands). Voltage offsets were maintained below 20 mV. Electrode placement followed the 10/20 international standard (Klem et al., 1958; Oostenveld and Praamstra, 2001). Offline preprocessing was conducted using the Python MNE toolbox version 1.7 (Gramfort, 2014). The data were re-referenced to a common average and band-pass filtered with a high-pass filter at 0.1 Hz and a low-pass filter at 100 Hz to reduce slow drifts and high-frequency noise. Only the standard 64 scalp electrodes were retained for analysis, and data were aligned with stimulus onset markers extracted from the Status channel and matched to behavioural timing information. Epochs were extracted from -200 to 800ms relative to each stimulus onset and baseline-corrected using the -200 to 0ms pre-stimulus interval. Finally, the data were downsampled to 200 Hz to reduce file size and computational load.

### 2.4 Pairwise Exemplar Decoding

To assess fine-grained visual discrimination of individual exemplars within each category, we applied multivariate pattern analysis (MVPA) to each timepoint in a time-resolved exemplar decoding analysis (Grootswagers et al., 2017). This approach tested whether neural activity patterns could be reliably distinguished between individual exemplars within a given category (e.g., Dog 1 *vs.* Dog 2 *vs.* Dog 3…), providing a measure of within-category representational specificity that is robust to low-level visual feature differences arising across different images of the same exemplar (Ventura et al., 2026). As each exemplar was represented by multiple naturally varying images differing in viewpoint, illumination, and other image properties. Because classifiers were trained and tested across these image variations, successful decoding could not rely on idiosyncratic low-level features of any single image. Instead, above-chance performance indicates that neural activity captured information that remained consistent across different appearances of the same exemplar. For each participant, and for each category separately (faces, cars, houses, dogs), we conducted a pairwise binary classification analysis between all possible exemplar pairs. At each time point, a Linear Discriminant Analysis (LDA) classifier was trained to discriminate between two exemplars based on the multivariate EEG pattern across all electrodes, using a sliding estimator approach, whereby classification was performed independently at each time point. Classification was implemented using a leave-one-block-out cross-validation scheme, ensuring independence between training and test data and controlling for potential block-level dependencies. For each fold, the classifier was trained on epochs sourced from all but one block and subsequently tested on epochs sourced from the held-out block. This procedure was repeated until each block had served as the test set.

Decoding accuracy was computed for each exemplar pair and averaged across all pairs within a category, yielding a time-resolved measure of exemplar discriminability. Chance performance for this binary classification was 50%. This analysis allowed us to quantify the extent to which individual exemplars could be distinguished within each category and to test whether Super-recognisers exhibit enhanced within-category discrimination relative to typical recognisers.

### 2.5 Representational Similarity Analysis

To characterise the structure of neural representations of individual exemplars both within and across categories, we used the pairwise decoding accuracies to construct time-resolved representational dissimilarity matrices (RDMs) for each participant and time point (Grootswagers et al., 2017). Because higher decoding accuracy indicates greater separability in the neural responses evoked by two exemplars, these RDMs reflect the unfolding representational geometry associated with discriminating individual exemplars within a given category, as well as the separability of exemplars from different categories. To test whether neural representations of exemplars captured the categorical distinction between faces and non-face objects, we rank-correlated (Spearman) the neural RDM for each participant and timepoint with a binary theoretical RDM that captured face versus non-face category structure (exemplars belonging to the same category were assigned a value of 0, whereas face–non-face pairs were assigned a value of 1), yielding a time-resolved measure of model fit. Correlation values were subsequently averaged across participants within each group (Super-recognisers and typical recognisers) to examine group-level differences in representational organisation.

To assess whether neural representational structure was organized similarly across individuals, we directly compared RDMs across participants. Because each participant’s RDM series captured the degree to which their neural responses distinguished between all pairwise combinations of exemplars, correlating these matrices across individual serves to quantify representational overlap over time. At each time point, we used Spearman correlation to compare pairwise discriminability values from one participant with those of every other participant within the same group. This yielded one SR-SR and one TR-TR time-resolved inter-subject similarity matrix in which each cell reflects the degree to which two individuals distinguished between exemplars according to a similar relational pattern. High similarity values indicated that stimulus pairs strongly separated in one participant’s neural responses were also strongly separated in others’, whereas lower values reflected greater divergence in representational structure. To assess category-specific representational structure, we considered intersubject similarity separately within each category (faces, cars, houses, dogs) to yield a time-resolved index of within-group similarity for *i)* Super-recognisers and *ii)* typical recognisers, separately for each category. Higher within-group similarity therefore reflects more stable and shared representational geometry for a given category.

To further isolate the geometry of face representations, we conducted a follow-up analysis focusing on pairwise relationships involving faces. Starting from the neural representational dissimilarity matrices (RDMs), we restricted the analysis to pairs that included a face together with a non-face, while also excluding same categories comparisons (e.g., face-face). This yielded a reduced representational space capturing the relative positioning of face stimuli with respect to other categories. For each participant and time point, we extracted the relevant pairwise dissimilarity values. Intersubject similarity was then computed again by correlating these values across participants within each group using Spearman correlation, yielding time-resolved measures of the consistency with which faces are represented relative to other categories, representing a targeted index of face-specific distinctiveness.

### 2.6 Statistical analysis

Statistical inference combined Bayesian analyses for time-resolved EEG measures with frequentist analyses of behavioural accuracy. Bayesian analyses were used because the EEG measures were evaluated continuously across time, and Bayes Factors allow evidence to be quantified in favour of both the alternative and null hypotheses. This approach therefore provided a time-resolved estimate of whether above-chance neural decoding, positive model correlations, or intersubject similarity were supported by the data, and when evidence instead favoured the absence of an effect (Teichmann, 2022). We used the BayesFactor R package (Morey & Rouder, 2018) for all Bayesian analyses. Bayesian t-tests were implemented using the default Cauchy prior on standardised effect sizes (*r* = 0.707; Wetzels et al., 2011). Depending on the specific analysis and hypothesis, Bayesian tests were conducted either directionally (e.g., testing for above-chance decoding or greater effects in Super-recognisers relative to typical recognisers) or non-directionally using standard two-tailed comparisons. Bayes Factor (BF) values greater than 3 were interpreted as moderate evidence for the alternative hypothesis, values between 1 and 3 as anecdotal evidence for the alternative, values between 1/3 and 1 as anecdotal evidence for the null, and values smaller than 1/3 as moderate evidence for the null hypothesis. For time-resolved EEG measures, Bayesian tests were performed independently at each time point. For decoding analyses, one-sample Bayesian t-tests were used to assess whether decoding accuracy exceeded chance level, defined as 50% for pairwise exemplar decoding. Model correlations were computed between neural RDMs and the theoretical face/non-face model RDM, and intersubject similarity values were computed by correlating representational structure across participants. For these Spearman correlation measures, one-sample Bayesian t-tests were used to assess whether correlations differed from zero within each group. Between-group differences were assessed using independent-samples Bayesian t-tests comparing Super-recognisers and typical recognisers at each time point. For visualisation, Bayes Factors were expressed on a log10 scale.

We analysed the surprise memory test data by subjecting participants’ conditional mean accuracy to a mixed-design ANOVA with a between-subject factor of *Group* (Super-recognisers, typical recognisers) and within-subjects factors of *Category* (faces, cars, houses, dogs) and *Trial Type* (novel, seen).

## Results

### 2.7 Behavioural recognition memory performance

To test that participants had encoded the rapid image sequences presented in the main experiment, we first inspected behavioural performance on the short surprise recognition task completed at the end of the experiment. As can be seen in Figure 2, recognition accuracies in nearly all conditions were well above chance, suggesting that observers in both groups encoded meaningful information about individual entities presented during the main experiment. We observed a main effect of Group, *F*(1, 40) = 31.14, *p* < .001, with Super-Recognisers achieving higher overall accuracy than typical recognisers (SR: *M* = .75, *SD* = .19; TR: *M* = .60, *SD* = .22). There was also a significant main effect of Category, *F*(3, 120) = 47.64, *p* < .001, indicating that recognition performance differed across stimulus categories. Recognition accuracy was highest for faces (*M* = .78), followed by dogs (*M* = .70), cars (*M* = .65), and houses (*M* = .57). FDR-corrected pairwise comparisons revealed significantly higher accuracy for faces than cars and houses, and significantly higher accuracy for dogs than houses. No other pairwise differences reached significance. Finally, there was a significant main effect of Trial Type, *F*(1, 40) = 7.93, *p* = .008, with higher recognition accuracy for previously encountered images (*M* = .74, *SD* = .18) relative to novel instances of the same exemplars (*M* = .60, *SD* = .17). No significant interactions between factors were observed (all *ps* > .18).

**Figure 2.**
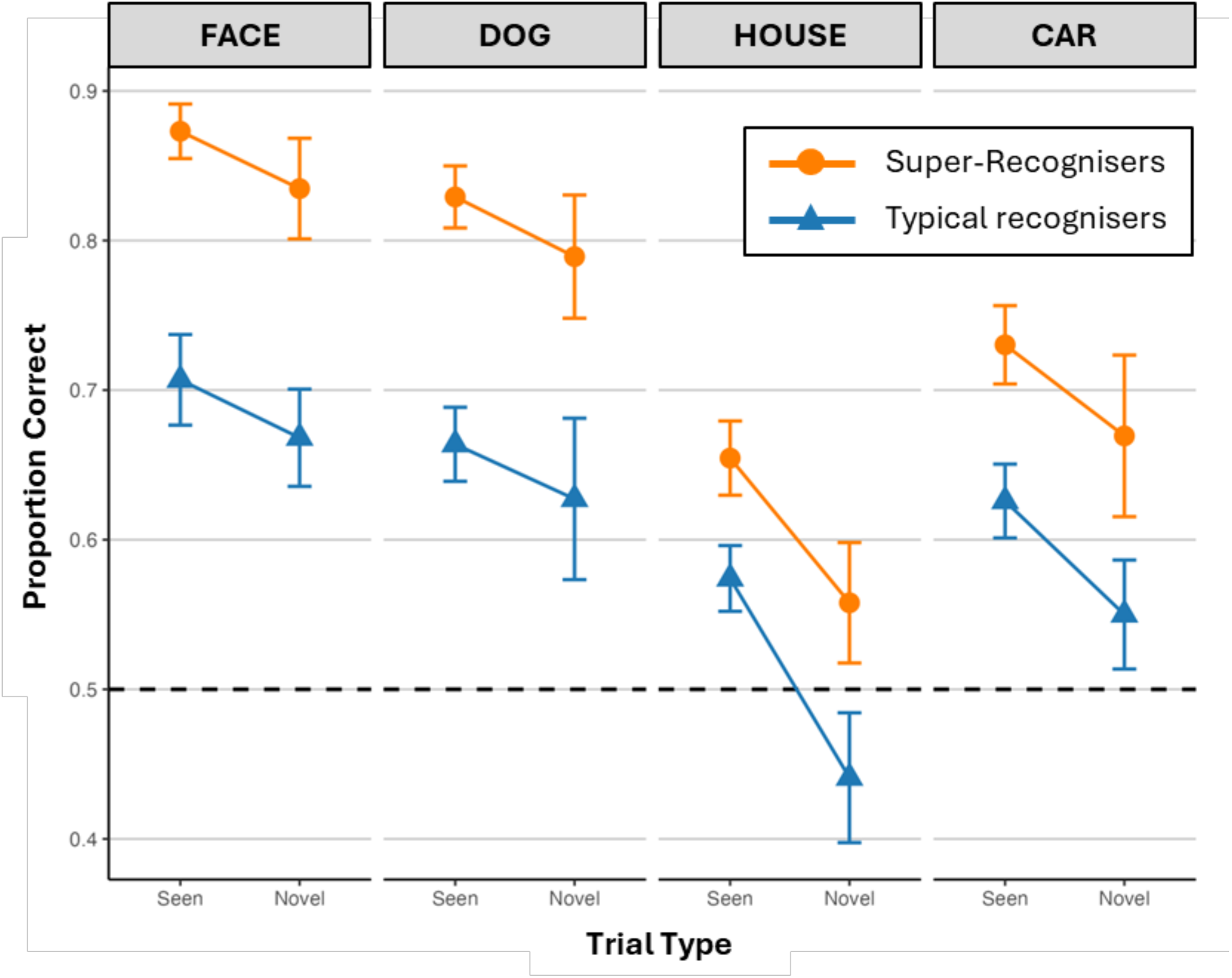
Surprise memory test behavioural performance as a function of *Group*, *Category* and *Trial Type*. Error bars represent ± SEM. Performance was well above chance (dashed line) for most conditions, suggesting observers encoded meaningful information about individual exemplars in all categories during rapid image sequences. Super-Recognisers showed higher overall recognition accuracy than typical recognisers across all stimulus categories. Across categories, recognition was higher for previously seen images than for novel images of previously encountered exemplars.

### 2.8 Neural Encoding of Individual Exemplars

Our key analysis sought to establish how well neural activity patterns in Super-Recognisers and typical recognisers distinguished between individual exemplars within each category. The graphs at the top of each panel of Figure 3 show mean pairwise decoding accuracy as a function of time, with panels A-D showing these data separately for faces, cars, houses, and dogs. Across all categories and both participant groups, decoding accuracy rose above chance shortly after stimulus onset, with reliable exemplar-level decoding emerging before approximately 100ms. However, the temporal profile of exemplar decoding differed across categories. Cars and houses exhibited relatively sharp early peaks in decoding accuracy emerging around ∼100-150ms and declining rapidly thereafter (Faces: BF_max_ = 1.2×10⁷ and 4.4×10⁴ for Super-recognisers and typical recognisers, respectively; Dogs: BF_max_ = 1.0×10⁷ and 3.4×10⁵; Houses: BF_max_ = 9.4×10⁶ and 6.7×10⁵; Cars: BF_max_ = 2.6×10⁵ and 1.2×10⁵.), consistent with processing dominated by lower-level visual feature differences. The dog category showed a somewhat more sustained temporal profile, with elevated decoding accuracy extending beyond the initial peak. Notably, face exemplar decoding exhibited the most temporally sustained profile across the epoch. Following an early rise comparable to the other categories, decoding of individual face identities remained elevated until approximately 700-800ms after stimulus onset, and this sustained identity information in later processing was observed mostly in Super-Recognisers.

**Figure 3.**
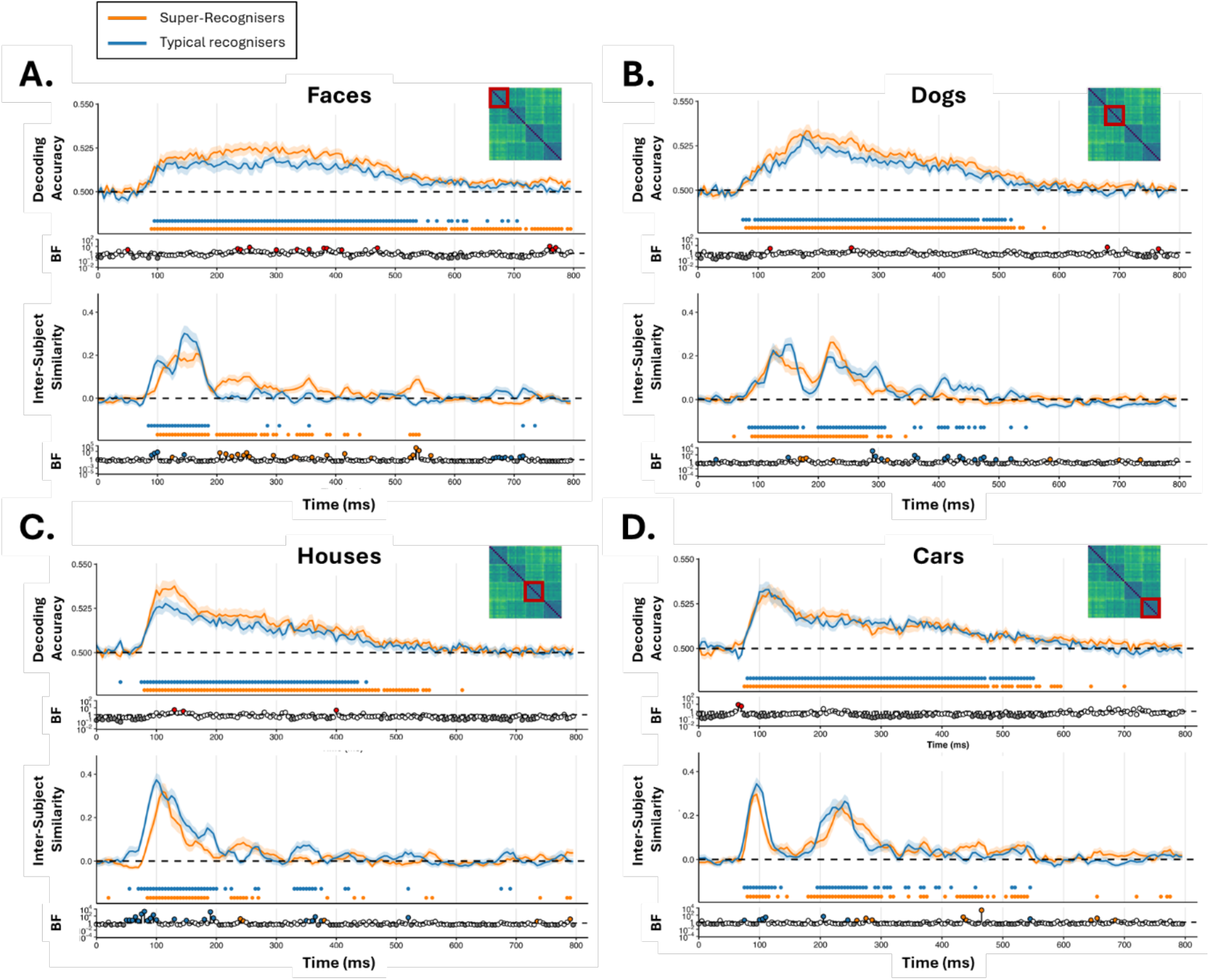
Time-resolved exemplar decoding accuracy and inter-subject representational similarity for (A) faces, (B) dogs, (C) houses, and (D) cars. Upper panels show mean pairwise exemplar decoding accuracy in each group, reflecting the ability of multivariate EEG activity patterns to distinguish between individual exemplars within each category (e.g., one face identity vs. another). Shaded regions are ± SEM. Lower panels show category-specific inter-subject representational similarity derived from representational dissimilarity matrices (RDMs) based on decoding accuracy and quantify the consistency of representational geometry across participants within each group. Insets illustrate the RDM submatrices corresponding to the stimulus pairs included in each analysis (highlighted in red). Dashed horizontal lines are chance performance (50%) for decoding and zero correlation for inter-subject similarity. Horizontal coloured markers below each plot indicate time points where there was moderate evidence (BF>3) for the relevant metric differing from baseline. The lowermost panel in each of A-D shows timepoint-wise Bayes Factors for a difference between the groups, with coloured points denoting moderate evidence (BF > 3) for one-tailed comparisons in decoding accuracy (SR>TR, red points) or two-tailed comparisons for inter-subject similarity (orange and blue points). Grey points denote evidence for the null (BF < 1/3).

Critically, group differences in exemplar decoding were not uniform across categories. Replicating our prior work (Ventura et al., 2026), we found that pairwise decoding of individual face identities was notably stronger in Super-recognisers’ neural responses than in typical recognisers’, particularly during the mid-latency interval (∼200-400ms). No other stimulus category demonstrated such group differences, with pairwise decoding profiles in Super-Recognisers and controls largely overlapping in the case of cars, houses, and dogs, with only small and transient differences observed over time. Thus, these findings suggest that Super-Recognisers’ perceptual encoding of exemplar-level differences was selectively enhanced for faces.

### 2.9 Inter-subject representational similarity

We next examined the consistency of representational geometry across individuals within each category using inter-subject representational similarity analysis (Ventura et al., 2026). Here, inter-subject similarity refers to the extent to which the relative organisation of exemplars within a category is shared across participants. As shown in the graphs at the bottom of each Figure 3 panel, inter-subject representational similarity emerged rapidly following stimulus onset for all categories in both Super-Recognisers and typical recognisers, beginning around ∼80-100ms, peaking between ∼120-200ms, and gradually declining at later latencies (>300ms). This temporal profile indicates that neural representations of individual exemplars became increasingly consistent across individuals during the early stages of visual processing before progressively weakening over time. Interestingly, faces exhibited the strongest and most sustained inter-subject similarity effects in both groups.

Critically, however, the temporal dynamics of face-related representational consistency differed substantially between Super-recognisers and typical recognisers. In Super-Recognisers, face representations remained reliably and consistently organised across participants from approximately ∼100ms until ∼350ms, forming a sustained period of stable representational organisation. In contrast, although typical recognisers also showed an early increase in face-related inter-subject similarity shortly after stimulus onset, this effect declined more rapidly at later latencies. Bayesian evidence for group differences was strongest during the ∼200-300ms interval, corresponding closely to the time window in which enhanced face exemplar decoding was also observed in Super-recognisers. For non-face categories (cars, houses, and dogs), both groups showed broadly comparable temporal profiles of inter-subject similarity, with little evidence for sustained group differences over time. These findings mirror the pattern observed above for pairwise decoding accuracy, in that group-level differences in inter-subject similarity were constrained to the face category. Thus, while neural representations of individual exemplars within each category became consistently organised across observers in both groups, Super-Recognisers exhibited a more sustained and stable shared representational structure than typical recognisers did, specifically in response to faces.

### 2.10 The Representational Boundary Between Faces and Objects

Having established differences in the organisation of neural representations for individual exemplars within each category, we next examined how faces were positioned relative to non-face categories within the broader representational structure. Here we began by assessing whether neural representations in each group reflected a face vs. non-face categorical structure using model-based representational similarity analysis. As shown in Figure 4A, correlations with the Face/Non-Face model increased rapidly following stimulus onset in both Super-recognisers and typical recognisers, emerging around ∼100ms and peaking between ∼120-200ms. This indicates that neural representations in both groups became increasingly organised according to a broad categorical distinction between faces and non-face objects.

**Figure 4.**
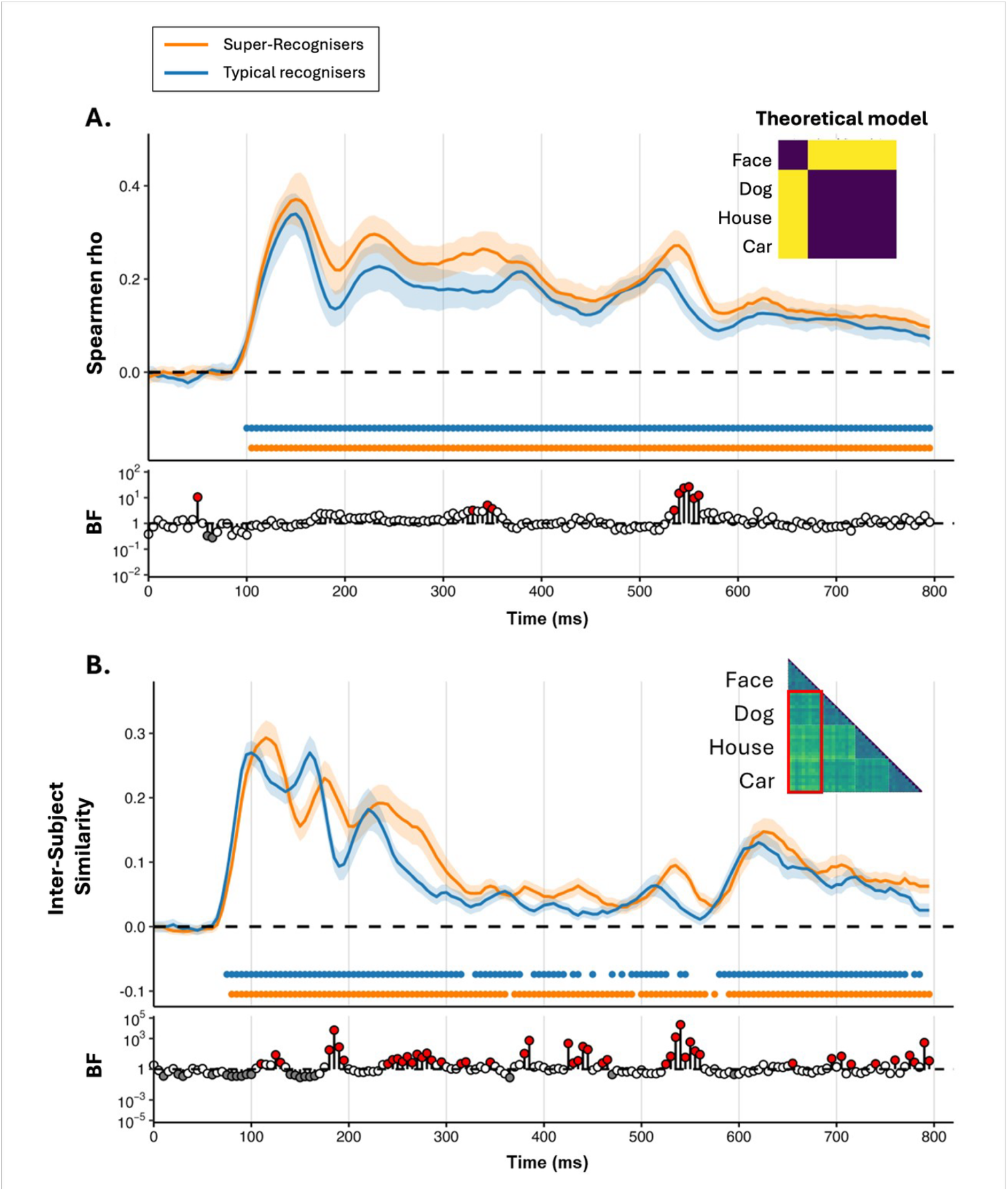
Time-resolved representational structure of faces relative to non-face categories. **(A)** Correlation between Super-Recognisers’ and typical viewers’ neural representational dissimilarity matrices (RDMs) and a theoretical face vs. non-face model (inset), quantifying the extent to which neural representations were organized according to a broad distinction between faces and non-face objects. **(B)** Inter-subject representational similarity for Super-Recognisers and typical recognisers, restricted to face vs. non-face comparisons (red outline in the inset RDM). This quantifies between-subject consistency with which faces were represented relative to non-face exemplars, separately within each group. Solid lines represent group means for Super-recognisers (orange) and typical recognisers (blue), with shaded regions indicating ± SEM. Dashed horizontal lines indicate zero correlation. Horizontal coloured markers below chance within each panel indicate time points with moderate evidence (BF > 3) for positive effects in each group. Lower panels show time-resolved Bayes Factors for the between-group comparison at each timepoint. Red points indicate moderate evidence (BF > 3) for greater effects in Super-recognisers than controls based on one-tailed Bayesian comparisons; grey points indicate evidence for the null (BF < 1/3).

Our prediction was that the representational separability of face and non-face categories would be stronger for Super-recognisers than typical recognisers, given their well-established behavioural advantages in face processing (e.g. Russel et al., 2009, Dunn et al., 2023). We found modest evidence for this possibility: Super-recognisers showed numerically higher model correlation values than typical recognisers, with some intervals showing moderate evidence favouring a group difference (BF > 3). Together, these results suggest that observers in both groups exhibited reliable face vs. non-face organisation of representational space, with modest evidence that this distinction may be comparatively stronger in Super-Recognisers than typical recognisers at a late stage of processing.

To further examine how faces were positioned relative to non-face categories, we conducted a targeted second-order representational similarity analysis (RSA) focusing specifically on representational consistency between individuals in terms of the face–non-face distinction. Rather than assessing the degree of inter-subject similarity reflected in the overall RDM structure (as in Figure 3), this analysis quantified inter-subject similarity specifically within the subset of image pairs that captured how faces are represented relative to non-face exemplars. For both groups, inter-subject similarity for the face–non-face representational separation increased soon after stimulus onset, with values rising around ∼90-100ms and peaking around ∼120-180ms. This indicates that the representational geometry that differentiated faces from non-face exemplars became increasingly consistent between individual observers in both groups over time. Notably, however, this representational consistency was stronger between individual Super-Recognisers than between individual control participants. Bayesian evidence favouring increased inter-subject similarity in face/non-face separation for Super-Recognisers relative to controls emerged during the early peak (∼120-180ms and 240-300ms) and again during later intervals (∼420-470ms and 520-560ms; one-tailed Bayesian comparison, BF > 3). Together, these findings suggest that faces were not only more consistently individuated in Super-Recognisers, but that they were more strongly separated from non-face exemplars within a broader representational structure.

## 3. Discussion

Here we tested whether individual differences in face recognition ability influenced neural encoding and differentiation of individual exemplars within various visual categories (faces, dogs, houses, cars). Our findings show clear support that Super-Recognisers exhibit selectively enhanced neural encoding of individual face identities, but not individual exemplars drawn from non-face categories. Neural discrimination of individual face exemplars was stronger and more sustained in Super-Recognisers than in typical-recogniser controls, with superior face recognition ability also associated with more consistent representational organisation of face information across individual participants.

### 3.1 The Neural Precision of Visual Expertise: Enhanced Encoding for Faces

Using time-resolved MVPA to examine the perceptual encoding of individual exemplars in different visual categories, we show that neural responses in both participant groups contain information separating individual exemplars within all visual categories from approximately 100ms onwards. Importantly, this analysis goes beyond standard category-level decoding (i.e., face *versus* dog *versus* house, etc). Rather than testing whether neural activity distinguished broad categories such as faces, cars, houses, and dogs, we asked whether neural responses differentiated individual exemplars contained within each category. Moreover, our use of naturalistic images meant that decoding identity required neural responses to generalise across substantial variation in the appearance of a given exemplar while also distinguishing that exemplar from others in the same category.

Notably, while individual exemplars within all four categories were reliably decodable in both typical recognisers and Super-Recognisers, the latter group demonstrated a sustained advantage in the neural encoding of individual face identities. Specifically, face exemplar decoding was reliably stronger in Super-Recognisers than controls, particularly in the mid latency range (∼200-300ms). However, comparable group differences were not observed for cars, houses, or dogs, where exemplar decoding profiles were broadly overlapping between Super-recognisers and typical recognisers. Crucially, our secondary inter-subject similarity analysis showed that this representational structure for face identities was also *more consistent* between individual Super-Recognisers than between individual control participants, suggesting that facial identity information is organized in a more stable and convergent manner in this group of high-performing individuals than it is in the typical brain (Ventura et al., 2026). These findings suggest that the neural advantage observed in Super-Recognisers lies in the ability to establish more distinct and stable representations of individual face identities during perceptual encoding. Faces place unusually high demands on fine-grained individuation because they share a common global configuration while differing in subtle identity-defining information (Ross et al., 2014; Lee, 2021). Our finding that this advantage was confined to faces, despite identical decoding procedures across all categories, therefore supports the view that exceptional face recognition reflects enhanced mechanisms for resolving highly similar facial identities rather than a general improvement in exemplar-level visual discrimination.

Indeed, we have recently shown a clear distinction between Super-Recognisers and typical one in terms of exemplar-level neural encoding of facial identity in Super-Recognisers (Ventura et al., 2026). The present findings replicate and extend this pattern, with Super-Recognisers again showing more selective, differentiated and temporally sustained neural separation of individual face identities.

This finding contrasts with other recent studies that have highlighted broader group-level differences extend beyond faces, where neural responses to diverse visual stimuli have been shown to discriminate between Super-Recognisers and typical viewers (Faghel-Soubeyrand et al., 2024; Nador et al., 2025). Neither of these prior studies reported group-level differences in face-selective or identity-selective neural activity, perhaps suggesting that the neural mechanisms underlying Super-Recognisers’ superior recognition capabilities are broadly tuned to heterogeneous object classes. However, these studies did not directly test for robust face identity neural signals in response to natural variation across multiple images of a face. By probing identity information in neural responses in this way, our findings provide evidence of enhanced neural differentiation of faces identity in the super-recogniser brain

To extract stable identity information from naturally variable images, the visual system must distinguish image-specific variability (e.g., viewpoint, illumination, and expression) from the more stable properties that differentiate one identity from another (Chang et al., 2021). While the initial feed-forward sweep of visual processing (∼100ms) is sufficient for broad category detection, this fine-grained individuation is thought to rely on subsequent recurrent processing to resolve visual ambiguity (Xie et al., 2025). Through natural visual experience, the brain learns to exploit how objects smoothly transform in the real world (Rolls, 2021), allowing neural representations to achieve a “temporal stability” that sustains a reliable identity code despite fleeting visual noise (Piasini et al., 2021).

The timing of the Super-Recogniser advantage, observed here during a mid-latency window (∼200-400 ms), provides some clues to the neural processing responsible for their superior abilities. Rather than reflecting early feed-forward sensory processing alone, this interval aligns more closely with stages associated with perceptual consolidation and representational refinement. This interpretation is broadly consistent with recent event-related potential research demonstrating that the emergence of stable, viewpoint-invariant facial identity representations maps onto this same temporal interval (e.g., the N250 component; ∼220-270ms), reflecting top-down processes involved in constructing progressively more abstract identity representations (Ventura et al., 2026). This window also coincides with other neural signatures associated with superior face recognition abilities reported elsewhere. For example, high-performing face recognisers exhibit enhanced mid-latency N250 responses during facial individuation, particularly for more distinctive identities within a multidimensional representational space (Schroeger et al., 2023), and we have recently found enhanced identity encoding in an overlapping mid-latency window in Super-Recognisers.

### 3.2 From Faces to Objects: The Structure of Visual Space

A clear representational boundary separating faces from non-face categories emerged rapidly from an early point in visual processing, peaking between 120-200ms in both typical recognisers and Super-Recognisers. This rapid categorical segregation aligns with contemporary accounts of ventral visual processing suggesting that object representations are organised across multiple representational levels, spanning low-level visual features, mid-level structural properties, and increasingly abstract category and semantic information (Clarke et al., 2013; Papale et al., 2019; Bracci & Op de Beek, 2023). Within this framework, the emergence of a face/non-face boundary reflects the progressive organisation of visual information into behaviourally meaningful object representations. While the face vs. non-face distinction was highly robust in both observer groups in our study, here we found that the representational separation of faces and non-faces was notably stronger in Super-Recognisers than in typical recognisers across several time windows (∼120-180ms; ∼240-300ms; ∼420-470ms; ∼520-560m).

The representational geometry of high-level visual cortex is known to shape how perceptual similarities are organised and experienced (Charest et al., 2014). Thus, actively stabilising diagnostic facial features and expanding the representational distances between highly overlapping facial identities may progressively reorganise the surrounding geometry of visual space itself. As facial exemplars become increasingly differentiated and stabilised, the entire face category becomes more strongly separated from neighbouring non-face object representations. Importantly, this amplified categorical boundary in Super-Recognisers is unlikely to reflect a distinct mechanism for broad category segregation. Rather, it may emerge naturally from the hyper-refined within-category individuation processes described above. This account is broadly consistent with evidence that object representations in inferotemporal cortex are organised within a continuous “object space” according to visual similarity rather than discrete category boundaries (Bao et al., 2020). From this perspective, enhanced within-face individuation would be expected to shift the position of faces within this broader representational space, increasing their separation from neighbouring object categories without requiring an independent mechanism for category segregation.

This pattern is broadly consistent with recent electrophysiological evidence suggesting that high performing face recognisers may possess a more differentiated and finely tuned multidimensional face-space (Schroeger et al., 2023). Accommodating the fine-grained representational distances required to individuate highly similar facial identities likely depends on increasing the relative separation between overlapping exemplars within this representational structure. As a consequence, stronger within-face differentiation may also contribute to a greater representational distinction between faces and non-face object categories. This suggests that the amplified categorical boundary observed in Super-Recognisers co-occurs with enhanced face exemplar discrimination, indicating that both effects may reflect a common enhancement of facial representations rather than separate mechanisms for identity and category processing.

### 3.3 From encoding to memory: Different Expressions of Super-recognisers’ Ability

Although Super-Recognisers in our study exhibited selectively enhanced neural encoding for faces, they also showed superior recognition memory performance across all stimulus categories in the surprise memory task, consistent with broader behavioural advantages reported elsewhere (Stehr et al., 2026). This pattern suggests that memory performance should not be viewed as a direct readout of the initial perceptual representation. Rather, perceptual representations are thought to undergo progressive transformation as information is integrated with downstream mnemonic processes, becoming increasingly abstract across the cortical hierarchy (Rademaker et al., 2019; Favila et al., 2022). Consequently, the face-specific encoding advantage observed here may reflect the quality of the initial perceptual code, whereas broader behavioural advantages may also depend on later memory-related computations supported by interactions between ventral visual and medial temporal lobe systems (Martin & Barense, 2023).

### 3.4 Conclusion

Evidence suggests that Super-Recognisers object processing abilities extend beyond the face domain (e.g., Dunn et al. 2023; Stehr et al., 2026), and some recent evidence suggests that neural responses when viewing non-face objects discriminates super-recognisers from typical viewers (Faghel-Soubeyrand et al., 2014). Our study shows clear evidence that superior face recognition ability is grounded in a selective enhancement of perceptual encoding mechanisms for faces, rather than a broad-based improvement in exemplar discrimination across visual categories. Where Super-Recognisers show stronger, more sustained, and more consistently organised neural representations of individual face identities, the same advantage does not extend to encoding exemplars from analogous object classes (dogs, cars, or houses). This specificity suggests that the mechanisms underlying superior face recognition may at least partially on category-specific representations rather than general perceptual processes.

## Data availability

The data and analysis code of this study will be made publicly available upon publication through an online repository.

